# Establishment of bovine extraembryonic endoderm cells

**DOI:** 10.1101/2024.12.17.628911

**Authors:** Hao Ming, Giovanna N. Scatolin, Alejandro Ojeda, Zongliang Jiang

## Abstract

Understanding the mechanisms of hypoblast development and its role in the implantation is critical for improving farm animal reproduction, but it is hampered by the lack of research models. Here we report that a chemical cocktail (FGF4, BMP4, IL-6, XAV939, and A83-01) enables de novo derivation and long-term culture of bovine extraembryonic endoderm cells (bXENs). Transcriptomic and epigenomic analyses confirmed the identity of bXENs and revealed that they are resemble hypoblast lineages of early bovine peri-implantation embryos. We showed that bXENs help maintain the stemness of bovine ESCs and prevent them from differentiation. In the presence of a signaling cocktail sustaining bXENs, the growth and progression of epiblasts are also facilitated in the developing pre-implantation embryo. Furthermore, through 3D assembly of bXENs with bovine ESCs and TSCs, we developed an improved bovine blastocyst like structure (bovine blastoid) that resembles blastocyst. The bovine XENs and blastoids established in this study represent accessible *in vitro* models for understanding hypoblast development and improving reproductive efficiency in livestock species.

## Introduction

During the mammalian pre-implantation development, the first lineage differentiation specifies the inner cell mass (ICM) and trophectoderm (TE) in the blastocyst; the ICM further differentiates into epiblast and hypoblast (or primitive endoderm) in blastocyst. Subsequently, hypoblast or primitive endoderm gives rise to the yolk sac by implantation and is critical to support early conceptus development by producing a spectrum of serum proteins, generating early blood cells, and transporting nutrient from the uterus to the embryo (1–3). The development of hypoblast is a very conserved process although its developmental timing varies among mammalian species. In cattle, the hypoblast specifics in day 8 blastocysts, and differentiates into yolk sac during implantation around day 18-23. The involution of yolk sac occurs 40 days postfertilization which companies the formation of the placenta (4–6). Particularly, hypoblast undergoes dynamic lineage development, which is coordinate with a period of rapid growth and elongation of embryo from spheroid form at day 9-11, to elongated form at day 12- 14 to a filamentous form at day 16 till implantation (7), when majority of pregnancy loss occur (8–10). Proper hypoblast development and function are pivotal for the success of pregnancy, however, our knowledge of hypoblast development, particularly in ruminant species, is limited due to technical and logistic difficulties associated with *in vivo* experiments and a lack of manipulatable cell culture models. Furthermore, whether and how extraembryonic tissues support the development of pre-implantation epiblast remain largely unknown.

Extraembryonic endoderm cells (XENs) are established from primitive endoderm of early embryos and represent valuable tools for studying hypoblast lineage differentiation and function during embryogenesis (11, 12). To date, the XENs have been established in multiple species including mice (12), porcine (13, 14), monkey (15), and humans (16). Notably, signaling pathways inducing XENs vary extensively among different mammals. Mouse XENs can be captured from ESCs via retinoic acid and Activin-A (17), while human hypoblast has been induced from naïve pluripotent stem cells dependent of FGF signaling (18) as well as a chemical cocktail (BMPs, IL-6, FGF4, A83-01, XAV939, PDGF-AA, and retinoic acid) (19). In the domestic species, porcine XENs can be derived from blastocysts using either LIF/FGF2 or LCDM (LIF, CHIR99021, (S)-(+)-dimethindene maleate, and minocycline hydrochloride) condition (13, 14). Attempts to establish bovine XENs from blastocysts have also identified FGF2 as a facilitator, but the resultant cells could not be maintained in long-term culture with limited characterization (20, 21). Thus far, the authentic bovine XENs have not been established yet.

In this study, we discovered that a modified chemical cocktail (FGF4, BMP4, IL-6, XAV939, and A83-01) supports de novo derivation and long-term culture of bovine XENs. We then attempted to use bovine XENs model epiblast and hypoblast lineage crosstalk and found that bovine XENs promote growth and stemness of bovine embryonic stem cells (ESCs). This is further confirmed during pre-implantation embryo development by supplement signaling cocktail sustaining bovine XENs in *in vitro* embryo culture. We observed that the growth and progression of epiblasts are facilitated in the developing pre-implantation embryo. Finally, by assembling bovine XENs generated in this study with expanded potential stem cells (EPSCs) (22) and trophoblast stem cells (TSCs) (23), we generated an improved self-organized bovine blastocyst-like structure (blastoid) that is more resemble blastocyst compared to the two lineage (ESC and TSC) assembled blastoids (24).

## Results

### De novo derivation of bovine XENs from blastocysts

We have previously shown that a combination of four molecules (FGF2, Activin-A, LIF, and Chir99021) was able to efficiently convert SOX2^+^ extended pluripotent stem cells (EPSCs) into SOX17^+^ hypoblast cells in bovine (24). Therefore, we first adapted these four factors (4F- XENM) to derive bovine XENs from day 8 hatched IVF blastocysts. We observed XEN-like colonies’ outgrowth (Fig. S1A). Since robust hypoblast markers remain largely unknown in bovine, by mining a single cell RNA-seq dataset of bovine day 12 embryo (7), we identified a group of novel bovine hypoblast marker genes including CTSV, FETUB, APOA1, APOE, COL4A1, and FN1 (**Fig. S1B, C**). We found that these bXEN-like cells highly expressed all identified bovine hypoblast markers, while barely expressed epiblast (SOX2, OCT4, and NANOG) or trophoblast (CDX2, GATA3, and GATA2) markers (**Fig. S1D**). However, these XENs can only be maintained up to ten passages, therefore, named as short-term passaged bovine XENs, or bXEN^S^.

Recently, human authentic hypoblast cells were successfully induced from naïve pluripotent stem cells with seven chemical molecules, including BMPs (a pSMAD1/5/9 activator), IL-6 (a pSTAT3 activator), FGF4, A83-01 (a pSMAD2 inhibitor and ALK4/5/7 inhibitor) and XAV939 (a WNT/β-catenin inhibitor and tankyrase inhibitor) along with PDGF-AA and retinoic acid (19). To further establish long-term culture of bovine XENs, we assessed the different combinations of these seven factors with the 4F-XENM. Our comprehensive screening process showed that neither seven factors nor adding individual or any combinations of seven factors into 4F-XENM medium could establish stable bovine XENs (**Table. S1**). Surprisingly, we found that a combination of FGF4, BMP4, IL-6, XAV939, and A83-01 (5F-XENM) was able to efficiently support the outgrowth of bXEN-like morphological colonies from blastocysts (**Fig. 1A**).

**Figure 1.**
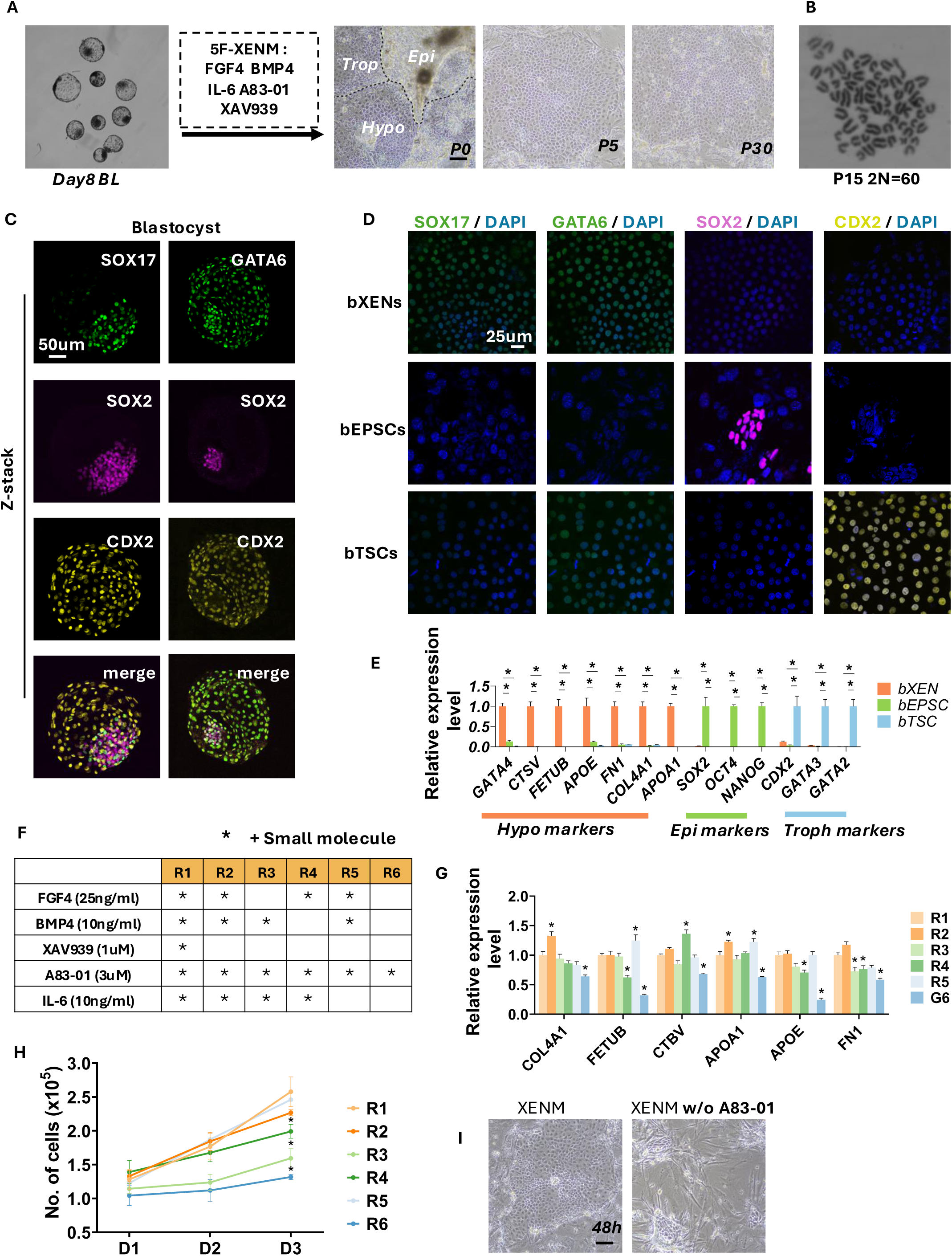
Derivation and characterization of bXENs. **A**. Representative image of outgrowth that formed from day 8 bovine blastocyst contains three morphologically distinct cell types and subsequent derivation and maintenance of bXENs. **B**. Karyotype analysis of bXENs at passage 15. **C**. Immunofluorescence analysis of GATA6 and SOX17 (hypoblast markers), SOX2 (epiblast marker), as well as CDX2 (trophectoderm marker) in day 8 blastocysts. Scale bar, 50μm. **D**. Immunofluorescence analysis of GATA6, SOX17, SOX2, as well as CDX2 in bXENs, bEPSCs, and bTSCs, separately. Scale bar, 25μm. **E**. Relative expression of defined lineage marker genes in three stem cell lines. **F.** The small molecules included in each medium recipe (R1-R6). **J.** Relative expression level of hypoblast marker genes in bXENs cultured in different mediums from **F**. **H.** Cell number estimated within 3 days following passage. **I.** Morphology of bXENs cultured in XENM with or without A83-01. Data are presented as the mean ± SEM. ∗*P* < 0.05 from one-way analysis of variance (ANOVA) followed by Tukey’s multiple comparisons test.

The derived bXENs-like colonies maintained stable and self-renewal properties with long-term passages (>30) (**Fig. 1A**), therefore named as long-term passaged bovine XENs, or bXENs. Further characterization revealed that bXENs maintained stable epithelial morphology of flattened colonies with clearly defined margins (**Fig. 1A**) and a normal diploid number of chromosomes (2N = 60) after long-term *in vitro* culture (**Fig. 1B**). Immunostaining analysis showed that, similar to hypoblasts of bovine blastocysts, bXENs expressed SOX17 and GATA6, but not either SOX2 or CDX2 which is positive in bEPSCs or bTSCs, respectively (**Fig. 1C, D**). They also highly expressed all identified novel hypoblast markers (**Fig. 1E**).

Furthermore, we determined the essential small molecules that are required for the maintenance of bXENs. First, A83-01 alone could maintain the stable expansion of bXENs while the cell proliferation and the marker gene expression were reduced (**Fig. 1F, G, H**). Withdrawal of A83-01 from 5F-XENM, bXENs exhibited differentiation and failed to expand into stable cell lines (**Fig. 1I**), suggesting A83-01 is indispensable in maintaining bXENs. Second, we observed that withdrawal of XAV939 or XAV939 together with IL-6 has a limited impact on the expression of bXENs’ marker genes (**Fig. 1G**) and cell proliferation (**Fig. 1H**). Third, absent of XAV939 and BMP4 resulted in reduced expression level of hypoblast markers such as FETUB, APOE, and FN1 (**Fig. 1G**). Finally, we demonstrated that BMP4 and FGF4 were two major factors affecting bXENs’ proliferation (**Fig. 1H**). Together, we demonstrated that 5F-XENM was effective in supporting the maintenance of bXENs.

Collectively, we developed a bovine XENs culture condition supports de novo derivation and self-renewal of stable bXENs *in vitro*.

## Transcriptional and chromatin accessibility features of bXENs

We next explored the transcriptomes of bXENs by RNA sequencing (RNA-seq) analysis. We compared the transcriptomes of bXENs with bovine EPSCs (22) and bovine TSCs (25). Principal component analysis (PCA) revealed that bXENs were clustered distinct from bEPSCs in PC1 and bTSCs in PC2, respectively (**Fig. 2A**), indicating the unique identity of three types of bovine stem cells, which is also shown in the correlation analysis (**Fig. S2A**). The identity of bXENs was further confirmed by the expression of representative marker genes of their corresponding blastocyst lineages (PrE, Epi, and TE) in these three types of bovine stem cells (**Fig. 2B**). Of note, bXENs also highly expressed both extraembryonic visceral endoderm (VE) and parietal endoderm (PE) markers (**Fig. 2B**), suggesting their developmental capacity towards VE and PE of the yolk sac. Additionally, we identified signaling pathways that were uniquely enriched in bXENs, including PI3K-Akt, cholesterol metabolism, focal adhesion, TNF, and TGF-beta signaling pathways (**Fig. 2C**). Intriguingly, when integrating transcriptomes of bXENs with single cell transcriptomes of hypoblast lineages of bovine peri-implantation embryos from day 12 through day 18 (7), we found that bXENs are closely clustered with highly proliferating hypoblasts from spheroid embryos (D12 and D14) while distinct from more differentiated hypoblasts of elongated embryos (D16 and D18), suggesting bXENs resemble early hypoblast populations *in vivo* (**Fig. 2D**).

**Figure 2.**
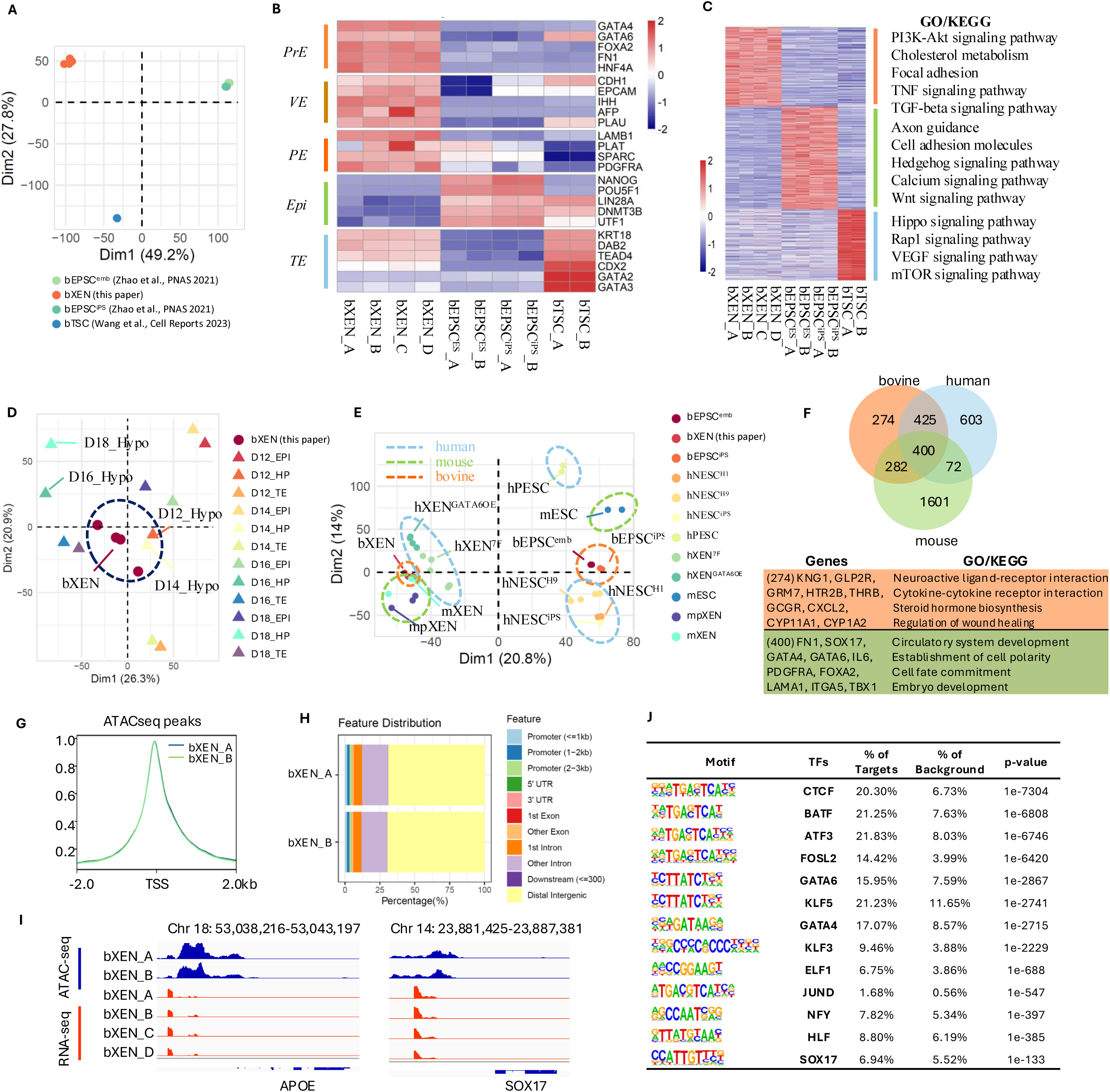
Transcriptomic and chromatin accessible features of bovine XENs. **A**. Principal component analysis (PCA) of transcriptomes of bXEN, bEPSC^emb^, bEPSC^iPS^, and bTSC. **B.** Heatmap showing the marker gene expression for PrE, VE, PE, Epi, and TE from each bovine stem cell type. **C.** Heatmap (left panel) showing lineage-specific expressed genes for bXENs, bEPSCs, and bTSCs, as well as the enriched signaling pathways (right panel). **D.** PCA of transcriptomes of bXENs and three major lineages from day 12-18 bovine *in vivo* embryos. **E.** PCA of transcriptomes of XEN and ESC from bovine, mouse, and human. Datasets of bEPSC^emb^ and bEPSC^iPS^ are from Zhao et al., PNAS 2021; Datasets of hNESC^H1^, hNESC^H9^, hNESC^iPS^, hPESC, hXEN^7F^, and hXEN^GATA6OE^ are from Okubo et al., Nature 2024. Datasets of mESC, mpXEN and mXEN are from Zhong et al., Stem Cell Research 2018. **F.** Venn diagram (top panel) showing the number of XEN enriched genes when compared to ESC among three mammalian species, as well as the enriched GO/KEGG (bottom panel) categories from overlapped genes and bovine specific genes, respectively. **G.** The enrichment of ATACseq peaks at annotated promoters (TSS ± 2kb) (normalized and on average) in bXENs. **H.** Feature distribution of ATACseq peaks in bXENs. **I.** The genome browser views showing the ATAC-seq peaks and RNA-seq reads enrichment near *APOE* and *SOX17* in bXENs. **F.** Motif enrichment analysis of ATAC-seq peaks from bXENs.

Additional transcriptomic comparisons of XENs and ESCs among cattle (22), human (19), and mice (26) further confirmed the lineage identity of bXENs (**Fig. 2E**). To further investigate the unique transcriptomic features of bXENs, we explored XEN specific genes compared to ESC in three mammalian species separately and identified 400 genes that are commonly enriched in XENs (**Fig. S2B**). These genes were involved in regulating canonical hypoblast functions, including circulatory system development, cell fate commitment, and embryo development (**Fig. 2F**). Additionally, 274 genes are uniquely in bovine, mainly manipulating ligand-receptor interaction, cytokine-cytokine receptor interaction, and steroid hormone biosynthesis (**Fig. 2F**). It is also noteworthy that bovine and human XENs share more common genes than those compared to mouse.

We also conducted ATAC-seq analysis to characterize the genome-wide chromatin accessibility of bXENs (**Fig. 2G, H**). Our analysis showed that the chromatin accessibilities of hypoblast lineage marker genes are consistent with their expression profiles (e.g., *APOE* and *SOX17*) (**Fig. 2I**). We confirmed that canonical hypoblast transcriptional factors (TFs) binding motifs are enriched in bXENs, such as *GATA6*, *GATA4, SOX17* (**Fig. 2J**). In addition, we identified several novel bovine hypoblast TFs including *CTCF, BATF, ATF3, FOSL2, KLF5, KLF3, ELF1, JUND*, and *NFY*, and *HLF* (**Fig. 2J**). We further validated these novel hypoblast TFs and confirmed that their chromatin accessibilities are consistent with their gene expression as well (**Fig. S2C**). Most of them (*CTCF, NFYA, NFYC, JUND*) were also highly expressed in the hypoblast cells of a day 12 peri-implantation bovine embryo (**Fig. S2D**).

Together, the RNA-seq and ATAC-seq analyses confirmed the molecular identity of bXENs and shed light on the molecular features during the earliest steps of hypoblast development in bovine.

## bXENs regulate the development of peri-implantation epiblasts

In cattle, the attachment of blastocysts is preceded by a period of rapid growth and elongation, when hypoblast lineages present dynamic changes between spheroid (D12 and D14) and elongated (D16 and D18) embryos, and have intense communication with both epiblast and trophectoderm lineages (7, 27). Given that both epiblast and hypoblast specify from ICM and the plasticity of two lineages are largely unknown, here we implemented a 3D co-culture model with our established robust bXENs to examine whether or how hypoblast regulates the development of epiblast during bovine embryogenesis. We mixed different ratio of bEPSC: bXEN cell populations (Group1 (G1): bEPSCs / bXENs = 40/0; Group2 (G2): bEPSCs / bXENs = 10/30; Group3 (G3): bEPSCs / bXENs = 10/0; and Group4 (G4): bEPSCs / bXENs = 0/40) (**Fig. 3A-D**). We found that co-cultures in both G1 and G2 can form spherical structures, but not those from G3 and G4 (**Fig. 3A-D**). Next, we conducted immunofluorescence analysis of SOX2 and GATA6 that are exclusively expressed in bEPSCs and bXENs, respectively. We found that spherical structures organized from bEPSCs alone in G1 largely remain SOX2 positive with GATA6 positive cells located at the peripheral region (**Fig. 3E**). Further quantification showed that aggregates in G1 consisted of three types of structures, including T1) SOX2^-^ GATA6^+^, T2) SOX2^+^ GATA6^-^, T3) SOX2^+^ GATA6^+^ (**Fig. 3E**). These results indicate a loss of pluripotency and randomly differentiation of EPSCs to hypoblasts-like cells in consistent with previous observations both in humans (19, 28) and bovine (24). On the contrary, co-culture of bEPSCs with bXENs in G2 resulted in spherical structures with cleaner and smoother periphery region compared to those of G1 (**Fig. 3B**), suggesting an improved survival of aggregates. Further immunostaining analysis showed that all cells in G2 remain SOX2 positive without any detection of GATA6^+^ cells (**Fig. 3F**), suggesting that the present of bXENs prevents bEPSCs from differentiation. To rule out the possibility that XENs transdifferentiate into SOX2^+^ cells, we tagged bXENs with GFP, followed by co-culture. We observed that bXENs were aggregated with bEPSCs on day 1, and all GFP^+^ bXENs disappeared by day 4 (**Fig. 3G**). Additionally, we found that, at the present of bXENs in G2, bEPSCs’ proliferation was significantly facilitated than those of G1, based on the size of formed spherical structures (**Fig. 3H**). These results demonstrated that the presence of bXENs and associated communications support the growth and stemness of bEPSCs.

**Figure 3.**
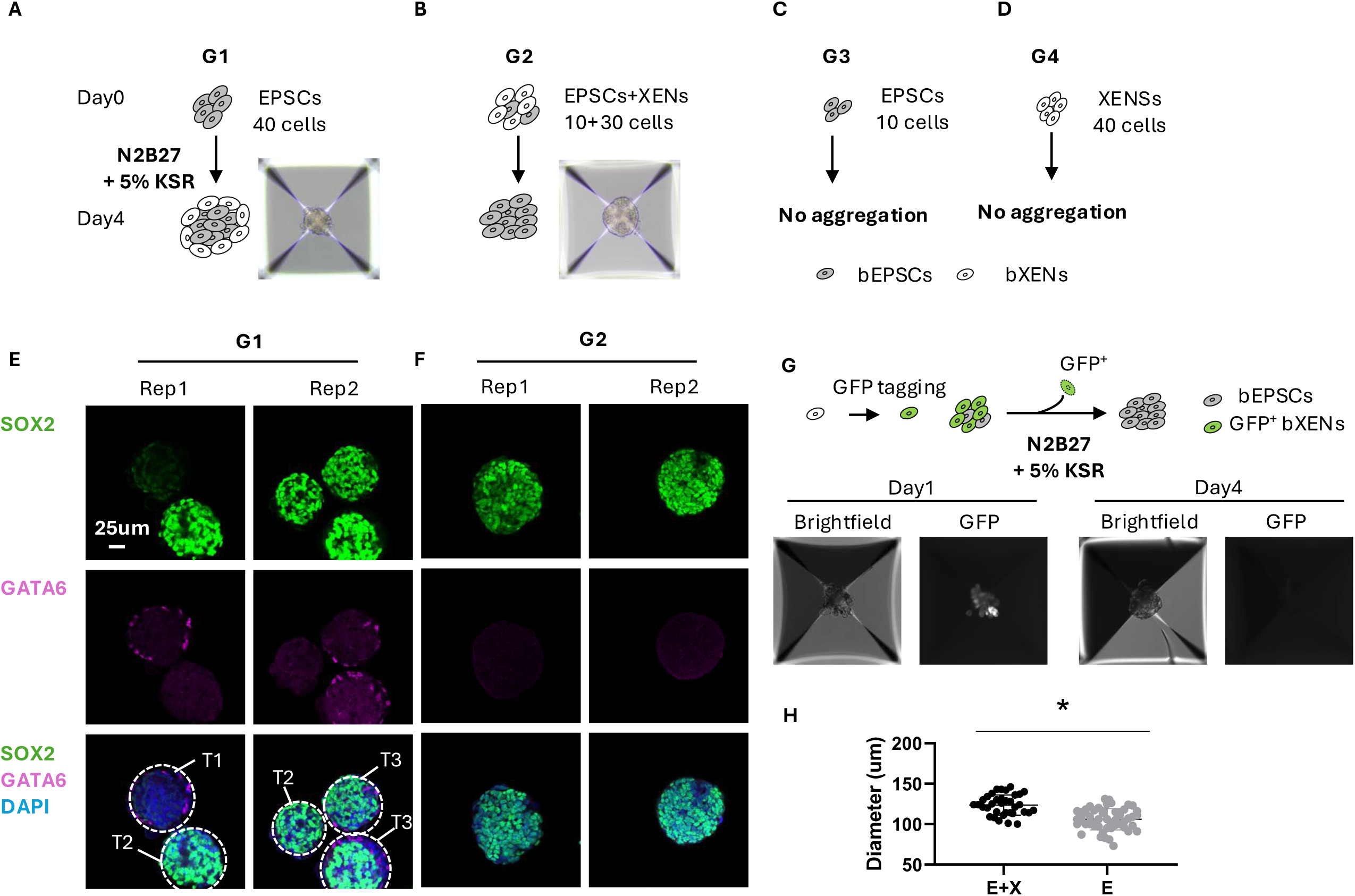
3D co-culture of bEPSCs and bXENs. **A.** Aggregates formed by 40 bEPSC cells cultured in N2B27 medium with 5% KSR. **B.** Aggregates formed by the mixture of bEPSCs and bXENs cultured in N2B27 medium with 5% KSR. **C.** Aggregates formed by 10 bEPSC cells cultured in N2B27 medium with 5% KSR. **D.** Aggregates formed by 40 bXEN cells cultured in N2B27 medium with 5% KSR. **E.** Immunofluorescence analysis of cell aggregates formed by bEPSCs. Green, SOX2; Pink, GATA6; Blue, DAPI. **F.** Immunofluorescence analysis of cell aggregates formed by mixture of bEPSCs and bXENs. Green, SOX2; Pink, GATA6; Blue, DAPI. **G.** Aggregates generated by the mixture of 10 bEPSCs and 30 GFP tagged bXENs on day1 and day4, respectively. **H.** Diameters of aggregates formed by the mixture of bEPSCs and bXENs (E+X) or by the bEPSCs (E) alone. Data are presented as the mean ± SEM. ∗*P* < 0.05 from unpaired *t*-test.

To further confirm the role of bXENs in promoting epiblast development, we treated bovine *in vitro* cultured embryos with defined small molecular cocktails sustaining bXENs (BMP4, FGF4, A83-01, XAV939, IL-6). The treatment was given at different developmental period before and after major genome activation or hypoblast specification (Experiment 1 (Exp. 1): day 1-8; Experiment 2 (Exp. 2): day 5-8; Experiment 3 (Exp. 3): day 8-12) (**Fig. 4A**). The subsequent developmental rate and lineage composition and allocation were measured by immunostaining analysis of epiblast marker SOX2, hypoblast marker SOX17, and trophectoderm marker CDX2. When treating embryos with bXEN signaling cocktails from either day 1-8 (Exp. 1) or day 5-8 (Exp. 2), we found day 8 hatched blastocysts had a significantly increased SOX2^+^ / SOX17^+^ cells ratio compared to the control group (**Fig. 4B, C**). We also noticed that these bXEN small molecules had no impact on early cleavage and trophoblast differentiation until blastocyst (**Fig. 4D**). However, the blastocyst hatching rate decreased dramatically presumably due to the issue of lineage specification within ICM (**Fig. 4D**). Further treating blastocysts with bXEN signaling cocktails during extended culture period from day 8 (Exp. 3), we observed that day 12 embryos had a well-defined and condensed SOX2^+^ spot in the ICM region (**Fig. 4E**), consistent with *in vivo* D12 embryos (**Fig. S3**), while ICM structure went through degeneration in control group (**Fig. 4E**). When calculating the ratio of SOX2^+^ and SOX17^+^ cells, we observed a significantly higher SOX2^+^ cells and lower SOX17^+^ cells in treated group compared to control. These results demonstrated that bXEN signaling cocktails could effectively protect epiblast from differentiation or degeneration.

**Figure 4.**
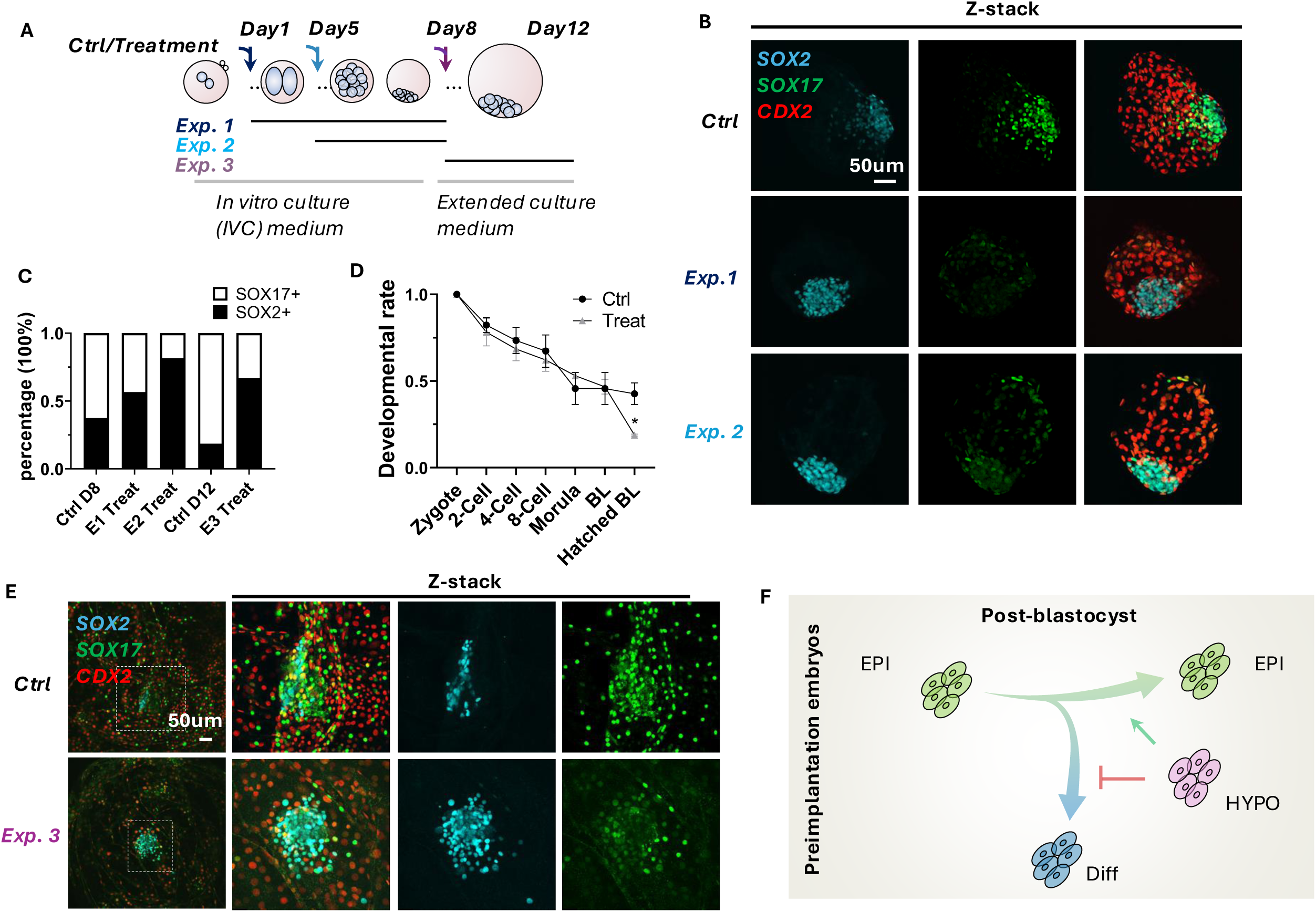
bXENs regulate the development of peri-implantation epiblasts. **A.** Schematic summarizing the treatment. The treatment was given at different developmental period before and after major genome activation or hypoblast specification (Experiment 1 (Exp. 1): day 1-8; Experiment 2 (Exp. 2): day 5-8; Experiment 3 (Exp. 3): day 8-12). **B.** Immunofluorescence analysis of SOX17 (hypoblast marker), SOX2 (epiblast marker), and CDX2 (trophectoderm) in day 8 blastocysts under the treatment in Exp.1 and Exp. 2. Scale bar, 50 μm. **C.** Ratio of SOX17^+^ and SOX2^+^ cells in embryos from Exp. 1-3. **D.** Developmental rates of embryos under Control or treatments. **E.** Immunofluorescence analysis of SOX17, SOX2, and CDX2 in day 12 embryos under the treatment in Exp. 3. Scale bar, 25 μm. **F.** Schematic summarizing the impact of hypoblast on maintenance or differentiation of epiblast. Data are presented as the mean ± SEM. ∗P < 0.05 from unpaired t-test.

Taking together, our experiments with both co-culture cell model and IVF embryos demonstrated that hypoblast regulate the development of epiblast in bovine by preventing its differentiation during bovine peri-implantation (**Fig. 4F**).

## Generation of bovine blastocyst-like structures by self-organization of bXENs, bEPSCs, and bTSCs

We have previously reported the successful generation of bovine blastoids by self-assembly of bEPSCs and bTSCs in tFACL+PD culture condition (24), which providing an accessible *in vitro* cell model for studying embryogenesis. However, the blastoids assembled by the two-lineage approach has shown lower proportion of hypoblast lineage compared to IVF blastocysts, which may limit their developmental capacity. The availability of bXENs prompted us to develop improved bovine blastoids through 3D assembly of bXENs, bEPSCs and bTSCs. We first aggregate three bovine stem cell types (bEPSC / bXEN / bTSC = 8:8:16) with the same culture condition (FGF2, Activin-A, Chir99021, Lif, and PD0325901) we reported (24). We found that this condition can support the formation of blastoids with high efficiency (46.60% ± 3.80%) within 4 days. The resulted blastoids contains a blastocele-like cavity, an outer TE-like layer, and an ICM-like compartment, morphologically equivalent to day 8 blastocysts (**Fig. 5A, D, E**). However, these blastoids have vanished SOX17^+^ hypoblasts compared to day 8 IVF blastocysts (**Fig. 5A, D**), same as we observed in our two-lineage assembled blastoids (2L-blastoids) (**Fig. 5C**) (24).

**Figure 5.**
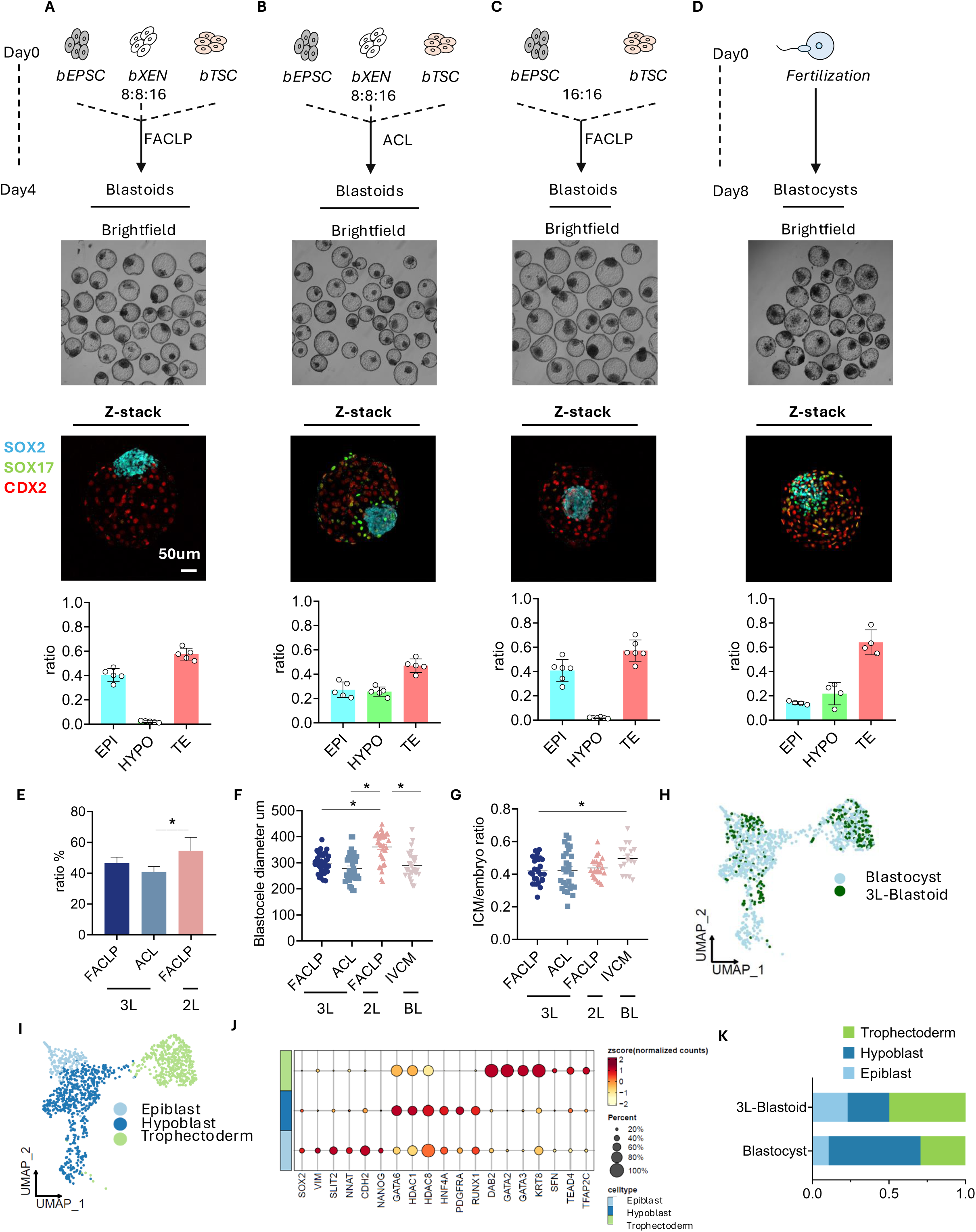
Generation of bovine blastocyst-like structures by self-organization of bXENs, bEPSCs, and bTSCs. A-D. Top panel: Illustration of the bovine blastoid formation using different assembly approach (**A**: bEPSCs, bTSCs, and bXENs aggregation in FACLP medium; **B**: bEPSCs, bTSCs, and bXENs aggregation in ACL medium; **C**: bEPSCs and bTSCs aggregation in FACLP medium as our previous published, as well as the IVF blastocysts control. Bottom panel: Phase-contrast and immunofluorescence analysis of blastoids from distinct protocols and blastocysts, as well as the quantification of lineage composition. Green, SOX17; Blue, SOX2; Red, CDX2. **E.** Blastoid foration rate from distinct protocols. **F.** Blastocele diameter measurement of blastoids from distinct protocols and blastocysts. **G.** Inner cell mass (ICM)/embryo ratio measurement of blastoids from distinct protocols, as well as blastocysts. **H.** Joint uniform manifold approximation and projection (UMAP) embedding of 10X Genomics single-nucleus transcriptomes of bovine 3L-blastoids (green) and bovine day 8 blastocyst (blue). **I.** Major cluster classification based on marker expression. **J.** Dot plot indicating the expression of markers of epiblast, trophectoderm, and hypoblast. **K.** Percentage of three cell types in blastoid and blastocyst. Data are presented as the mean±SEM. ∗*P* < 0.05 from one-way analysis of variance (ANOVA) followed by Tukey’s multiple comparisons test.

It has been shown that FGF2 could bias the cell fate of ICM towards PrE (24, 29, 30). As we integrated XENs, the FGF2 is not necessary for the blastoid induction anymore. Also, MEK inhibitor PD0325901 inhibits hypoblast specification from ICM (29, 31), which might be the reason for the vanished hypoblasts. Therefore, we withdrawn both FGF2 and PD0325901 from the culture condition, and found that the modified medium, ACL (Activin-A, Chir99021, Lif) supported the formation of blastoids morphologically resemble day 8 IVF blastocysts (**Fig. 5B**). With this approach (3L-blastoids), the blastoid formation efficiency reached 40.84% ± 4.76% within 4 days. Importantly, 3L-blastoids had a similar proportion of hypoblast and a slightly higher ratio of epiblast population compared to day 8 blastocysts, with majority hypoblast cells surrounding epiblasts (**Fig. 5B, D**). Additionally, the blastocelle size and ICM/blastocelle ratio of 3L-blastoid were also equivalent to day-8 IVF blastocysts (**Fig. 5F, G**). Thus, the bovine blastoid established in this study were more closely resembled blastocysts compared to the 2L-blastoids (24).

To determine the transcriptional states of 3L-blastoids, we performed single-nucleus RNA sequencing (snRNAseq) analysis of 3L-blastoids using the 10x genomics low throughput (up to 1,000 cells) platform. To ensure the precisely comparation, we generate the first snRNA-seq dataset of bovine day 8 IVF blastocysts using the same 10x genomics platform. Joint uniform manifold approximation and projection (UMAP) analysis revealed overall cells from 3L- blastoid clustered well with blastocyst-derived cells (**Fig. 5H**). We annotated three major cell clusters from blastocysts representing three blastocyst lineages, including Cluster 1 highly expressed SOX2, VIM, SLIT2, NNAT, CDH2, and NANOG as epiblasts, Cluster 2 highly expressed GATA6, HDAC1, HDAC8, HNF4A, PDGFRA, and RUNX1 as hypoblast, and Cluster 3 highly expressed DAB2, GATA2/3, KRT8, SFN, TEAD4, TFAP2C as trophectoderm cells (**Fig. 5I, J**). Of note, in the blastocyst, the defined epiblast cells still expressed hypoblast markers, and vice versa (Fig. 5J), suggesting the segregation of epiblast and hypoblast within ICM has not completed yet (32). The marker gene expression had the same patterns in all three annotated lineages from both 3L-blastoids and blastocysts, such as GATA2 (TE markers), PDGFRA (HYPO markers), and SLIT2 (EPI markers), indicating 3L-blastoid transcriptionally resemble to blastocyst (**Fig. S4A**). The comparative clustering analysis of blastoid and blastocyst cells showed that blastoids have a lower hypoblast cell population and higher epiblast population compared to the blastocysts (**Fig. 5K**). This confounding factor in single cell gene expression analysis may constitute the difference of lineage composition from immunostaining analysis. Additionally, we performed GO analysis of genes specifically enriched in each of three lineages of blastoids. We found genes specific to epiblast involve in regulating nervous system development, cell junction organization, and stem cell population maintenance, genes upregulated in hypoblast regulate cell morphogenesis and cell fate commitment, and finally genes highly expressed in trophectoderm involve in manipulating lipid biosynthetic process, actin cytoskeleton organization, and cell migration (**Fig. S4B**). Intriguingly, it was shown that most of the lineage specific genes in epiblast and hypoblast lineages were transcription factors (TFs) or TF cofactors, unveiling the lineage specific functions of those critical TFs during lineage specification within ICM and the further differential events (**Fig. S4B**).

Together, we developed an efficient and robust protocol to generate bovine blastoids by assembling bXENs, bEPSCs, and bTSCs that can self-organize and faithfully recreate all blastocyst lineages.

## Discussion

Hypoblast and its derivatives play a vital role in supporting and patterning the embryo (33), however, owing to applicable approaches associated with *in vivo* experiments, knowledge of hypoblast lineage segregation and development remains largely unknown. Here we demonstrated that a chemical cocktail (FGF4, BMP4, IL-6, XAV939, and A83-01) could support de novo derivation and maintenance of stable bXENs from bovine IVF blastocysts. Hypoblast lineage segregation and development is a conserved progress, while signaling to specify and sustain hypoblast is divergence among mammalian species. In mice, XENs do not require FGF signaling and can be maintained in the presence of retinoic acid and Activin-A (17). In humans, hypoblast induction requires FGF signaling (18) and can also be induced from naïve pluripotent stem cells using a chemical cocktail (BMPs, IL-6, FGF4, A83-01, XAV939, PDGF-AA, and retinoic acid) (19). In the domestic species, porcine XENs can be derived from blastocysts using either LIF/FGF2 or LCDM (LIF, CHIR99021, (S)-(+)-dimethindene maleate, and minocycline hydrochloride) (13, 14). Of note, LCDM is reported to maintain both bovine iPSCs (34) and TSCs (25). Interestingly, previous studies have demonstrated that FGF2 is also a key factor maintaining bovine primitive endoderm cell cells (6, 21). In the presence of FGF2, bEPSCs could also efficiently produce XENs (24). Here we have also shown that bXENs cells can be induced in the presence of FGF2, however, they are only capable for short-term self-renewal. Instead, a modification of human hypoblast induction condition by removing retinoic acid supports long-term culture of bXENs. The bXENs established in this study fill a gap and add a reliable stem cell model for research into pre- and peri-implantation development of an ungulate species.

In this study, we have demonstrated the regulatory role of hypoblast in regulating epiblast development in a ruminant species. This has also been most recently highlighted in both mouse and humans using *in vitro* experimental models (19, 28, 35). In humans, the existence of hypoblast can facilitate ESCs in generating a pro-amniotic-like cavity, which recapitulates the anterior-posterior pattern and mimics several aspects of the post-implantation embryo (19, 28). In mouse, primitive endoderm stem cells supported the lineage plasticity and the PrE alone was sufficient to regenerate a complete blastocyst and continue post-implantation development. Unlike many other mammalian species, ruminant species undergo a unique conceptus elongation process before implantation. During this phase, dramatic proliferation and differentiation of trophoblast and hypoblast lineage occur while the germ layer differentiation from epiblast is only observed until day 16 (7), which may constitute limited developmental potential of bovine XENs compared to humans and mouse. Thus, the bovine XENs model established here have mirrored the physiological lineage interaction between hypoblast and epiblast during bovine conceptus elongation.

A final contribution of this study is the development of an improved protocol to assemble bovine blastoids by self-organization of bXENs, bEPSCs, and bTSCs. In our previous study, the 2 lineages (EPSC and TSC) assembled bovine blastoids had a lower proportion of hypoblast lineages compared to the IVF blastocysts (24), which may compromise their developmental potential. Instead, with the integration of bXENs, we have shown that the 3L-blastoids established here more closely resemble bovine blastocysts compared to the first generation of bovine blastoids in terms of morphology, lineage composition and allocation, and transcriptional features. So far, most of the established blastoid models from other species, especially human, are self-organized from naïve ESCs or EPSCs (36–38), which comes up with concern that the differentiated TE-like lineage is transcriptionally resemble to amniotic ectoderm (39). By utilizing the assembly approach with authentic TSCs could eliminate this concern. Strikingly, embryo-like structures were generated through assembling mouse ESCs, TSCs, and XENs *in vitro*, which recapitulates the developmental characteristics of mouse embryos up to day 8.5 (40, 41), demonstrating that blastoids derived by assembling approach with three lineages possess higher developmental potential. This is significant in large mammals, particularly for the domestic species, as blastoid technology established here, upon further optimization and *in vivo* function testing, could lead to the development of novel artificial reproductive technologies for cattle breeding, which may enable a paradigm shift in livestock reproduction.

In summary, our work has established an authentic extraembryonic endoderm cell line and developed an improved bovine blastoid technology. We have also shown the valuable of bovine XEN in modeling cell-cell communications, thus filling a significant knowledge gap in the study of bovine embryogenesis when most of pregnancy failure occurs.

## METHODS

### Key resources table

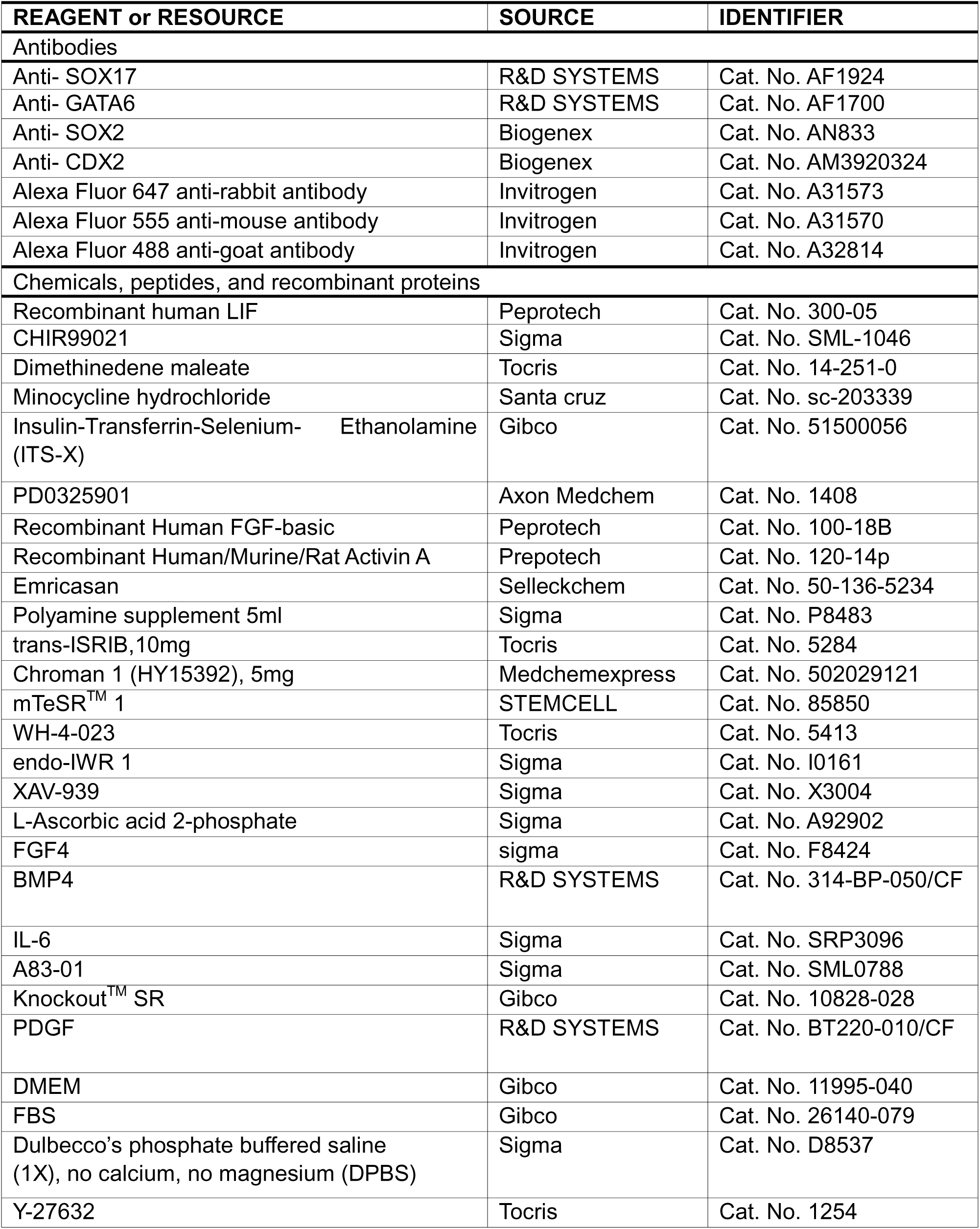
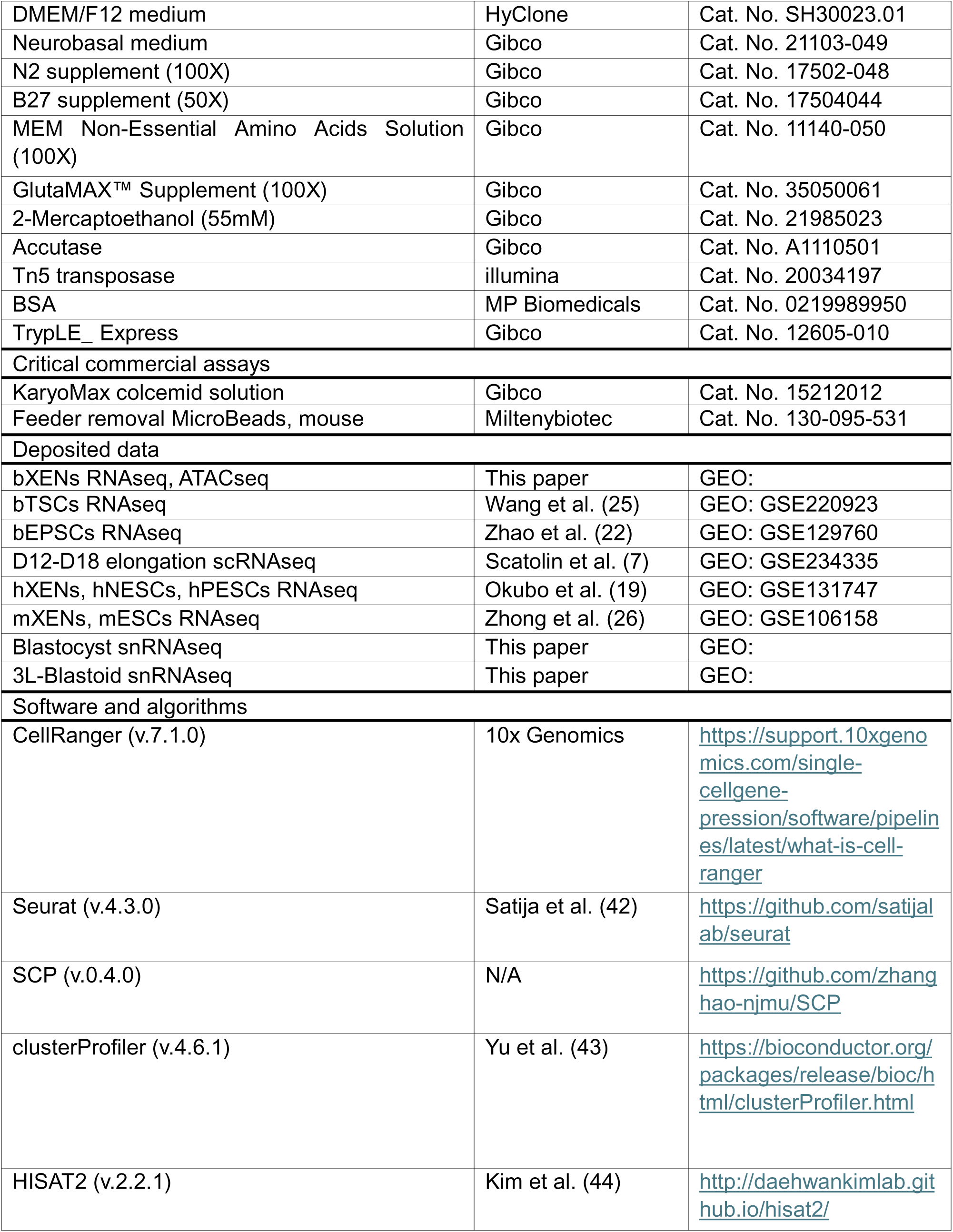
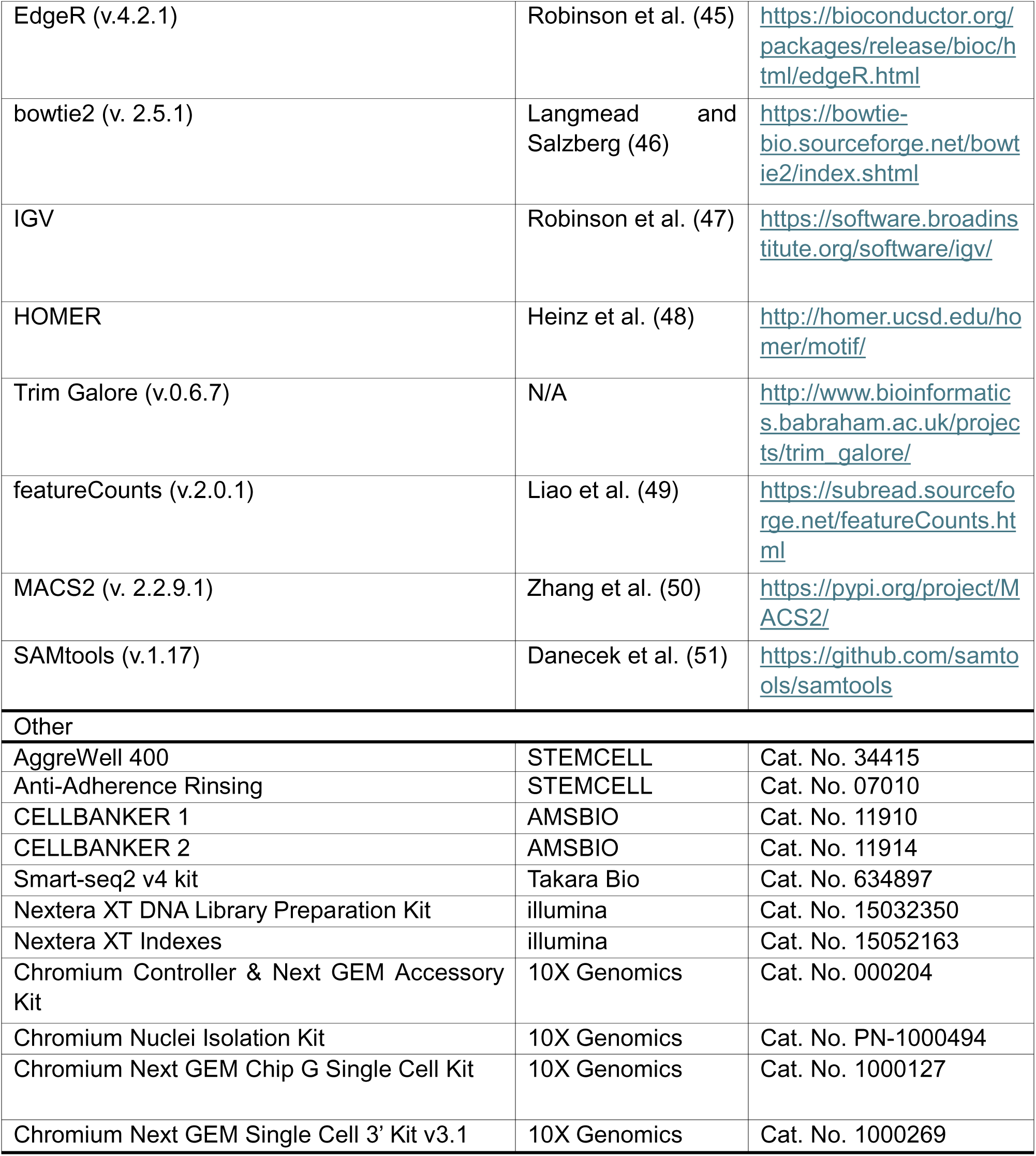

## RESOURCE AVAILABILITY

**Lead contact:** Further information for resources and reagents should be directed to the lead contact, Zongliang Jiang.

**Materials availability:** This study did not generate new unique reagents.

## Data and code availability

- The raw FASTQ files and normalized read accounts per gene are available at Gene Expression Omnibus (GEO) (https://www.ncbi.nlm.nih.gov/geo/) under the accession number GSE283042 and GSE283048. This paper analyzes publicly available data. The accession numbers for the datasets are listed in the key resources table.
- This paper does not report original code.
- Any additional information required to reanalyze the data reported in this paper is available from the lead contact upon request.

## METHOD DETAILS

### Bovine IVF embryo production

The IVF embryos used in this study were produced as previously described (52). Briefly, bovine cumulus-oocyte complexes (COCs) were aspirated from selected follicles of slaughterhouse ovaries. BO-IVM medium (IVF Bioscience) was used for oocyte *in vitro* maturation. IVF was performed using cryopreserved semen from a Holstein bull with proven fertility. Embryos were then washed and cultured in BO-IVC medium (IVF Bioscience) at 38.5 °C with 6% CO2. Day 8 hatched blastocysts were collected and were processed for bXENs derivation. For post-hatching culture from day8 until day12, embryos were transferred into extended culture medium containing DMEM: F12 (Gibco) and Neurobasal medium (Gibco) (1:1), 1x N2-supplement (Gibco), 1x B27-supplement (Gibco), 1x NEAA (Gibco), 1x GlutaMAX (Gibco), 0.1 mM 2-mercaptoethanol (Gibco), 10 μM/mL ROCK inhibitor (Tocris, 1254), 20 ng/mL ActivinA (Peprotech, 100-18B). For the treatment group, 25 ng/mL FGF4 (sigma, F8424), 10 ng/mL BMP4 (R&D SYSTEMS, 314-BP-050/CF), 1μM XAV939 (sigma, X3004-5MG), 3μM A83-01 (sigma SML0788-25MG), 10ng/mL IL-6 (sigma, SRP3096) were added.

### Derivation and culture of bXENs

ICMs were isolated from day 8 blastocysts by microsurgery and were placed in separate wells of a 24-well plate that was seeded with mitomycin C-treated mouse embryonic fibroblast (MEF) cells. To derive bXEN^S^, the ICMs were initially cultured in pre-bXENM^S^ containing DMEM: F12 and Neurobasal medium (1:1), 1x N2-supplement, 1x B27-supplement, 1x NEAA, 1x GlutaMAX, 0.1 mM 2-mercaptoethanol, 20 ng/mL bFGF (Peprotech, 100-18B), 20 ng/mL ActivinA. After three days, when all the ICMs attach and form outgrowth, change the medium to bXENM^S^ containing DMEM: F12 and Neurobasal medium (1:1), 0.5x N2-supplement, 0.5x B27- supplement, 0.5x NEAA, 0.5x GlutaMAX, 0.1 mM 2-mercaptoethanol, 1mM NaPy (Sigma, s8636), 10 μg/ml l-ascorbic acid (Sigma, A92902), 1x ITS-X (Gibco, 51500-056), 0.1% FBS, 0.5% KSR, 20 ng/mL LIF (Peprotech, 300-05), 1μM Chir99021 (Sigma, SML-1046), 10 ng/mL bFGF, 10 ng/mL ActivinA.

To derive bXEN, the ICMs were cultured in 5F-XENM containing DMEM: F12 and Neurobasal medium (1:1), 1x N2-supplement, 1x B27-supplement, 1x NEAA, 1x GlutaMAX, 0.1 mM 2-mercaptoethanol, 25 ng/mL FGF4, 10 ng/mL BMP4, 1μM XAV939, 3μM A83-01, 10ng/mL IL-6. To promote the proliferation of hypoblast, 0.1% KSR, 0.1% BSA (MP biomedicals), 20ng/mL ActivinA, and 10 ng/mL PDGF (R&D SYSTEMS, BT220-010/CF) were also added during the derivation of bXEN and were optional for bXENs maintenance. All the cells were cultured at 38.5°C and 5% CO2. The culture medium was changed every other day. On day 7 or 8, outgrowths were dissociated by TrypLE (Gibco, 12605-010) for 5-7 mins at 38.5 °C and passaged. For optimal survival rate, 10 μM Rho-associated protein kinase (ROCK) inhibitor Y- 27632 was added to the culture medium during first 24 hours. Once established, both bXEN^S^ and bXEN were passaged every 3-5 days at a 1:5 split ratio using TrypLE. Each well of cells was dissociated by 0.5mL TrypLE for 5 mins at 38.5°C, the same volume of DMEM/F12 with 1% BSA was used to neutralize. bXENs could be cryopreserved by CELLBANKER 2 (ZENOGEN) according to the manufacturer’s instructions.

### Bovine EPSCs culture

Bovine EPSCs were maintained in bovine EPSC culture medium (3i+LAF) (22): mTeSR base (STEMCELL, 85850), 1% BSA,10ng/ml LIF, 20ng/ml Activin A, 0.3 μM WH-4-023, 1 μM Chir99021, 5 μM IWRI, 50 μg/ml Ascorbic acid (Vitamin C). bEPSCs were passaged every 2 days at a 1:6 split ratio using TrypLE. Each well of cells was dissociated by 0.5mL TrypLE for 5 mins at 38.5°C, the same volume of bXEN medium was used to neutralize TrypLE, ROCK inhibitor is necessary for each passage. bEPSCs could be cryopreserved by CELLBANKER 2 according to the manufacturer’s instructions.

### Bovine TSCs culture

Bovine TSCs were derived and cultured in LCDM (25) (hLIF, CHIR99021, DiM and MiH) media (DMEM: F12 and Neurobasal medium (1:1), 0.5x N2-supplement, 0.5x B27-supplement, 1x NEAA, 1x GlutaMAX, 0.1 mM 2-mercaptoethanol, 0.1% BSA (MP biomedicals), 10 ng/mL LIF, 3 μM CHIR99021, 2 μM Dimethinedene maleate (DiM) (Tocris, 1425) and 2 μM Minocycline hydrochloride (MiH) (Santa cruz, sc-203339). bTSCs were passaged every 4 days at a 1:4 split ratio using Accutase (Gibco, A1110501). Each well of bTSCs was dissociated by 1mL Accutase for 5 mins at 38.5°C, the same volume of bTSCs medium was used to dilute Accutase for neutralizing the reaction. bTSCs were cryopreserved by CELLBANKER 1 according to the manufacturer’s instructions.

### Generation of reporter bXENs

The pLenti CMV GFP Puro plasmid (Addgene #17448) was packaged into lentivirus in 293T cells using JetPrime reagent (Polyplus, 101000015) following manufacturer’s instructions. After 48 hours incubation, the medium was collected and concentrated using the Lenti-X™ Concentrator (Takara, 631231). Then the GFP-lentiviruses were transfected into bXENs with 5μg/ml polybrene (sigma, TR-1003-G). 1μg/mL of puromycin (sigma, P8833) was added to the culture medium 2–3 days after transfection. Drug-resistant colonies with GFP signaling were manually picked and further expanded.

### Blastoid formation

For 3L-blastoid formation, bEPSCs, bTSCs, and bXENs were dissociated into single cells by treating with TrypLE for 3min, 15min, and 7min, respectively, followed by neutralizing with the same volume of their culture medium. After centrifugation at 300 x g for 5min, cells were resuspended in their normal culture media with ROCK inhibitor. Single-cell dissociation was made by gentle but constant pipetting until no visible cell clumps exist. To deplete iMEF cells, cells of three cell lines were filtered through passing 40mm cell strainers (Corning) separately and placed in precoated 6 well plates (Corning) with 0.1% gelatin and incubated for 35 minutes at 38.5 °C. Cells were then collected and stained with 1x trypan blue and manually counted in a Neubauer chamber. Current protocol is optimized for 8 bEPSCs, 8 bXENs and 16 bTSCs per well in a 1200 well Aggrewell 400 microwell culture plate (Stemcell technologies) for 9,600, 9,600, and 19,200 of each cell type per well. Each well was precoated with 500ml of Anti-Adherence Rinsing Solution (Stemcell technologies) and spun for 5 minutes at 1500 x g. Wells were rinsed with 1ml of PBS just before aggregation. The cells for one well were mixed and centrifuged at 300 x g for 5min, followed by resuspension with 1mL ACL medium (DMEM: F12 and Neurobasal medium (1:1), 1x N2-supplement, 1x B27-supplement, 1x NEAA, 1x GlutaMAX, 0.1 mM 2-mercaptoethanol, 0.5x ITS-X, 20 ng/ml LIF, 10 ng/ml ActivinA, 1μM Chir99021, supplemented with 1x CEPT cocktail (53) (50 nM chroman-1 (C, Tocris), 5 μM emricasan (E, Selleckchem), 0.7 μM trans-ISRIB (T, Tocris), and 1 x polyamine supplement (P, Thermo)). To ensure even distribution of the cells within each microwell, cells were gently mixed by pipetting with a P200 pipette. The plate was first placed in incubator for 8 min to allow the cells to settle down, then centrifuged at 700 x g for 3 minutes and put back to incubator. Half of the medium was changed daily, the blastoids showed up from day 2 and could be collected on day 3 or day4.

### Bovine EPSCs and XENs aggregation assay

bEPSCs, and bXENs were dissociated into single cells and depleted feeder cells as described above. Ten bEPSCs and 30 bXENs, or 40 bEPSCs, or 10 bEPSCs, or 40 bXENs seeded in each well of a 1200 well Aggrewell 400 microwell culture plate under N2B27 medium (DMEM: F12 and Neurobasal medium (1:1), 1x N2-supplement, 1x B27-supplement, 1x NEAA, 1x GlutaMAX, 0.1 mM 2-mercaptoethanol) with 5% KSR and 10 μM Y27632. Half of the medium was changed daily and the aggregation structures were cultured until day 4.

### Karyotyping assay

bXENs were incubated with 1 mL KaryoMAX colcemid solution (Gibco, 15212012) at 38.5°C for 4-5 hours. Cells were then dissociated using 1 mL Trypsin (Gibco, 25200-056) at 38.5°C and centrifuged at 300 x g for 5 min. The cells were resuspended in 1mL PBS solution and centrifuged at 400 x g for 2 min. The supernatant was aspirated and 500 mL 0.56% KCI was added to resuspend the cells. The cells were incubated for 15 min, then centrifuged at 400 x g for 2 min. 1 mL cold fresh Carnoy’s fixative (3:1 methanol: acetic acid) was added to resuspend the cells, followed by a 10 min incubation on ice. After centrifuge, 200 mL Carnoy’s fixative was added to resuspend the cells. Cells were dropped on the clean slides and air dried and soaked in a solution (1:25 of Giemsa stain (Sigma, GS500): deionized water) for 9 min.

Slides were rinsed with deionized water and air dried. The images were taken by Leica DM6B at 1000x magnification under oil immersion.

### Immunofluorescence analysis

Cells, blastoids, and blastocysts were fixed in 4% paraformaldehyde (PFA) for 20 min at room temperature, and then rinsed in wash buffer (0.1% Triton X-100 and 0.1% polyvinyl pyrrolidone in PBS) three times. Following fixation, cells were permeabilized with 1%Triton X- 100 in PBS for 30 min and then rinsed with wash buffer. Samples were then transferred to blocking buffer (0.1% Triton X-100, 1% BSA, 0.1 M glycine, 10% donkey serum) for 2 hours at room temperature. Subsequently, the cells were incubated with the primary antibodies overnight at 4°C. The primary antibodies used in this experiment include anti-SOX2 (Biogenex, an833), anti-CDX2 (Biogenex, MU392A; 1:100), anti-GATA6 (R&D SYSTEMS, AF1700; 1:100), and anti-SOX17 (R&D SYSTEMS, AF1924; 1:100). For secondary antibody incubation, the cells were incubated with Fluor 488- or 555- or 647-conjugated secondary antibodies for 1 hour at room temperature. Followed by DAPI staining (Invitrogen, D1306) for 15 min. The images were taken with a fluorescence confocal microscope (Leica).

### Quantitative real-time PCR

Total RNA was extracted from cells using RNeasy Micro Kit (Qiagen) according to the manufacture’s protocol. First-strand cDNA was synthesized using the iScript cDNA Synthesis Kit (BIO-RAD). The qRT-PCR was performed using SYBR Green PCR Master Mix (BIORAD) with specific primers (**Table. S2**). Data was analyzed using the BIO-RAD software provided with the instrument. The relative gene expression values were calculated using the ΔΔCT method and normalized to internal control beta-actin.

### RNA-seq library preparation and data analysis

Total RNA of bXENs was extracted using RNeasy Micro Kit (Qiagen). The RNA-seq libraries were generated using the Smart-seq2 v4 kit with minor modifications from the manufacturer’s instructions. Briefly, individual cells were lysed, and mRNA was captured and amplified with the Smart-seq2 v4 kit (Clontech). After AMPure XP beads purification, the high-quality amplified RNAs were subject to library preparation using Nextera XT DNA Library Preparation Kit (Illumina) and multiplexed by Nextera XT Indexes (Illumina). The concentration of sequencing libraries was determined using Qubit dsDNA HS Assay Kit (Life Technologies) and KAPA Library Quantification Kits (KAPA Biosystems). The size of sequencing libraries was determined using the Agilent D5000 ScreenTape with Tapestation 4200 system (Agilent). Pooled indexed libraries were then sequenced on the Illumina HiSeq X platform with 150-bp pair-end reads.

Multiplexed sequencing reads that passed filters were trimmed to remove low-quality reads and adaptors by Trim Galore (version 0.6.7) (-q 25 –length 20 –max_n 3 –stringency 3). The quality of reads after filtering was assessed by FastQC, followed by alignment to the bovine genome (ARS-UCD1.3) by HISAT2 (version 2.2.1) with default parameters. The output SAM files were converted to BAM files and sorted using SAMtools6 (version 1.14). Read counts of all samples were quantified using featureCounts (version 2.0.1) with the bovine genome as a reference and were adjusted to provide counts per million (CPM) mapped reads. Principal component analysis and cluster analysis were performed with R (a free software environment for statistical computing and graphics). Differentially expressed genes (DEGs) were identified using edgeR in R. Genes were considered differentially expressed when they provided a false discovery rate of <0.05 and fold change >2. ClusterProfiler was used to reveal the Gene Ontology and KEGG pathways in R.

### ATAC-seq library preparation and data analysis

The ATAC-seq libraries from fresh cells were prepared as previously described (52). Shortly, cells were lysed on ice, then incubated with the Tn5 transposase (TDE1, Illumina) and tagmentation buffer. Tagmentated DNA was purified using MinElute Reaction Cleanup Kit (Qiagen). The ATAC-seq libraries were amplified by Illumina TrueSeq primers and multiplexed by index primers. Finally, high quality indexed libraries were pooled together and sequenced on Illumina NovaSeq platform with 150-bp paired-end reads.

The ATACseq analysis followed our established analysis pipeline (52). All quality assessed ATAC-seq reads were aligned to the bovine reference genome using Bowtie 2.3 with following options: –very-sensitive -X 2000 –no-mixed –no-discordant. Alignments resulted from PCR duplicates or locations in mitochondria were excluded. Only unique alignments within each sample were retained for subsequent analysis. ATAC-seq peaks were called separately for each sample by MACS2 with following options: –keep-dup all –nolambda –nomodel. The ATAC-seq bigwig files were generated using bamcoverage from deeptools. The ATAC-seq signals were normalized in the Integrative Genome Viewer genome browser. The enrichment of transcriptional factor motifs in peaks was evaluated using HOMER (http://homer.ucsd.edu/homer/motif/).

### Single nuclei isolation and library preparation

The snRNA-seq libraries from frozen blastoids and day 8 blastocysts were prepared using Chromium Nuclei Isolation Kit (10x Genomics,PN-1000493) with minor modifications from the manufacturer’s instructions. In brief, frozen blastoids and day 8 blastocysts were transferred to pre-chilled sample dissociation tube and were dissociated with pestle within lysis buffer, then the tube was incubated on ice for 7 min. Then the dissociated sample was pipetted onto assembled nuclei isolation column and centrifuged 16,000 x rcf for 20 sec. After being washed with debris removal buffer and wash buffer, the nuclei pellet was resuspended in 50 μl resuspension buffer and performed cell counting. Nucleus were loaded into a 10x Genomics Chromium Chip following manufacturer instruction (10x Genomics, Chromium Next GEM Single Cel 3’ Reagent Kit v3.1 Dual Index) and sequenced with an Illumina Novaseq 6000 System (Novogene).

### snRNA-seq data analysis

To analyze 10X Genomics single-cell data, the base call files (BCL) were transferred to FASTQ files by using CellRanger (v.7.1.0) mkfastq with default parameters, followed by aligning to the most recent bovine reference genome downloaded from Ensembl database (UCD1.3), then the doublets were detected and removed from single cells by using Scrublet (0.2.3) with default parameters. The generated count matrices from all the samples were integrated by R package Seurat (4.3.0) utilizing canonical correlation analysis (CCA) with default parameters (https://satijalab.org/seurat/articles/get_started.html). The data was scaled for linear dimension reduction and non-linear reduction using principal component analysis (PCA) and UMAP, respectively. The following clustering and visualization were performed by using the Seurat standard workflow with the parameters ‘‘dim = 1:10’’ in ‘‘FindNeighbors’’ function and ‘‘resolution =0.5’’ in ‘‘FindClusters’’ function. The function ‘‘FindAllMarkers’’ in Seurat was used to identify differentially expressed genes in each defined cluster. The cutoff value to define the differentially expressed genes was p.adjust value <0.05, and fold change >0.25. The UMAP plots and bubble plots with marker genes were generated using ‘‘CellDimPlot’’ and ‘‘GroupHeatmap’’ functions in R package SCP (0.4.0) (https://github.com/zhanghao-njmu/SCP), respectively. Gene ontology (GO) and pathway analysis were performed using R package clusterProfiler (4.6.1).

### Quantification and statistical analysis

Data were analyzed using GraphPad Prism 9 (GraphPad Software, Inc.) unless otherwise stated. Two-tailed unpaired or paired *t*-tests were used to determine the significance of differences between the means of two groups. One-way ANOVA followed by multiple comparisons was used to determine the significance of differences among means of more than two groups. Values of p < 0.05 were considered statistically significant. Statistical analyses of sequencing data were performed in R. Genes with |fold change| >2 and p value < 0.05 were identified as significant DEGs. Gene ontology (GO) and pathway analysis were performed using R package clusterProfiler (4.6.1). The value of p value < 0.05 was considered significant.

## Author contributions

H.M, and Z.J designed the research. H.M conducted all research experiments and data analysis. G.S., and A. O. assisted with embryo experiments. Z.J supervised the research. H.M, and Z.J wrote the manuscripts with inputs from all co-authors.

## Acknowledgements

This work was supported by the NIH Eunice Kennedy Shriver National Institute of Child Health and Human Development (R01HD102533), the USDA National Institute of Food and Agriculture (2019-67016-29863), and Genus plc.

## Significance

Bovine embryo-derived stem cells hold the potential to substantially advance biotechnology and agriculture. Here, we report the derivation and long-term culture of bovine extraembryonic endoderm cells (XENs) from pre-implantation embryos. Importantly, this study not only demonstrates the utility of bovine XEN models in elucidating the mechanistic features of early bovine embryogenesis, but also develops an improved bovine blastocyst-like structures (blastoids) technology for the creation of novel assisted reproductive technologies.

## Supplementary Figures and Tables

**Figure S1.** Derivation and characterization of bXEN^S^ cells. **A**. Representative image of outgrowth that formed from day 8 bovine blastocyst and subsequent generations of bXEN^S^. **B.** UMAP analysis of transcriptomic dataset of day 12 *in vivo* embryos (Scatolin et al., iScience, 2024) revealing four distinct cell types identified as epiblast (EPI), hypoblast (HP), trophectoderm (TE), and intermediate (Int) cells. Dot plot representing the expression of gene markers for Epi, HP, and TE. Dot size represents the percentage of cells in the cluster expressing the gene markers, the color gradient represents the level of expression from high (red) to low (yellow). **C**. UMAPs analysis showing the expression levels of selected hypoblast markers (*CTSV, FETUB, APOA1, APOE, COL4A1, FN1*) in day 12 *in vivo* embryos among all clusters. The color gradient from gray to blue at the right refers to the gene expression level (high expression = blue). **D.** Relative expression of defined lineage marker genes in three cell lines. Data are presented as the mean±SEM. ∗*P* < 0.05 from one-way analysis of variance (ANOVA) followed by Tukey’s multiple comparisons test.

**Figure S2.** Transcriptional and epigenomic features of bovine XENs. **A.** Heatmap showing the correlation between bXEN, bEPSC, and bTSC. The color gradient represents the level of correlation from high (red) to low (blue). **B.** Heatmap showing overlapped XEN enriched genes among human, mouse, and bovine. **C.** The genome browser views showing the ATAC-seq peaks and RNA-seq reads enrichment near *NFYA, NFYC, CTCF* and *JUND* in bXENs. **D.** UMAPs showing the expression levels of specific transcription factors (*NFYA, NFYC, CTCF and JUND*) in day 12 *in vivo* embryos among all clusters. The color gradient from gray to purple at the right refers to the gene expression level (high expression = purple). Data are presented as the mean ± SEM. ∗*P* < 0.05 from one-way analysis of variance (ANOVA) followed by Tukey’s multiple comparisons test.

**Figure S3.** Immunofluorescence analysis of GATA6 (hypoblast), SOX2 (epiblast), as well as CDX2 (trophectoderm) in day 12 *in vivo* embryos. **Scale bar, 100μm.**

**Figure S4.** Single cell RNA-seq analysis of blastoid and blastocyst. **A.** UMAP showing expression of trophectoderm (GATA2), hypoblast (PDGFRA), and epiblast markers (SLIT2), respectively. **B.** Heatmap showing the expression of cell lineage-specific genes in epiblast, trophectoderm, and hypoblast, as well as the biological functions regulated by the genes, separately.

**Table S1.** Culture conditions screened for the derivation of bovine XENs.

