## Supplementary figures and images for "Establishment of bovine extraembryonic endoderm cells"

### Figure S1

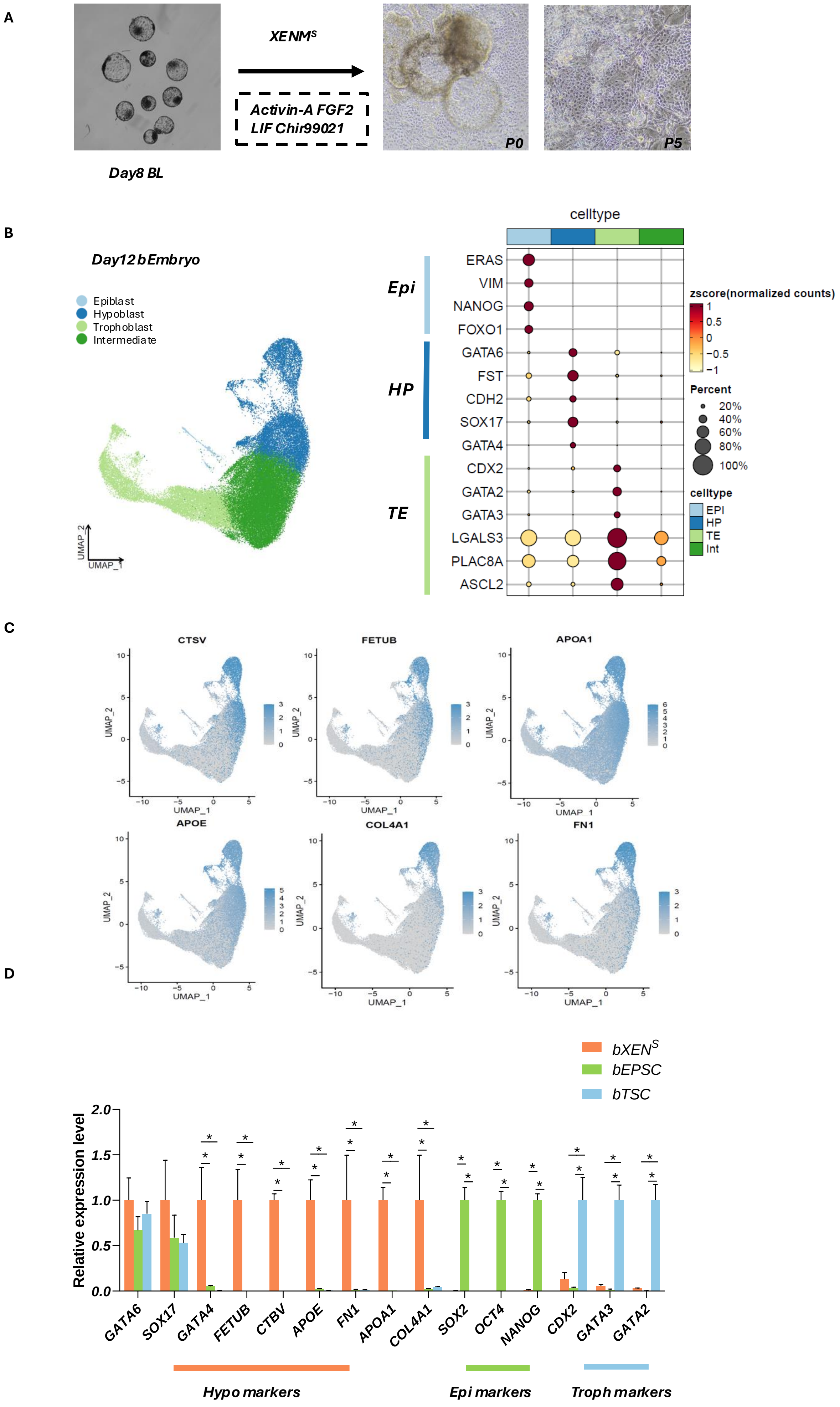

### Figure S2

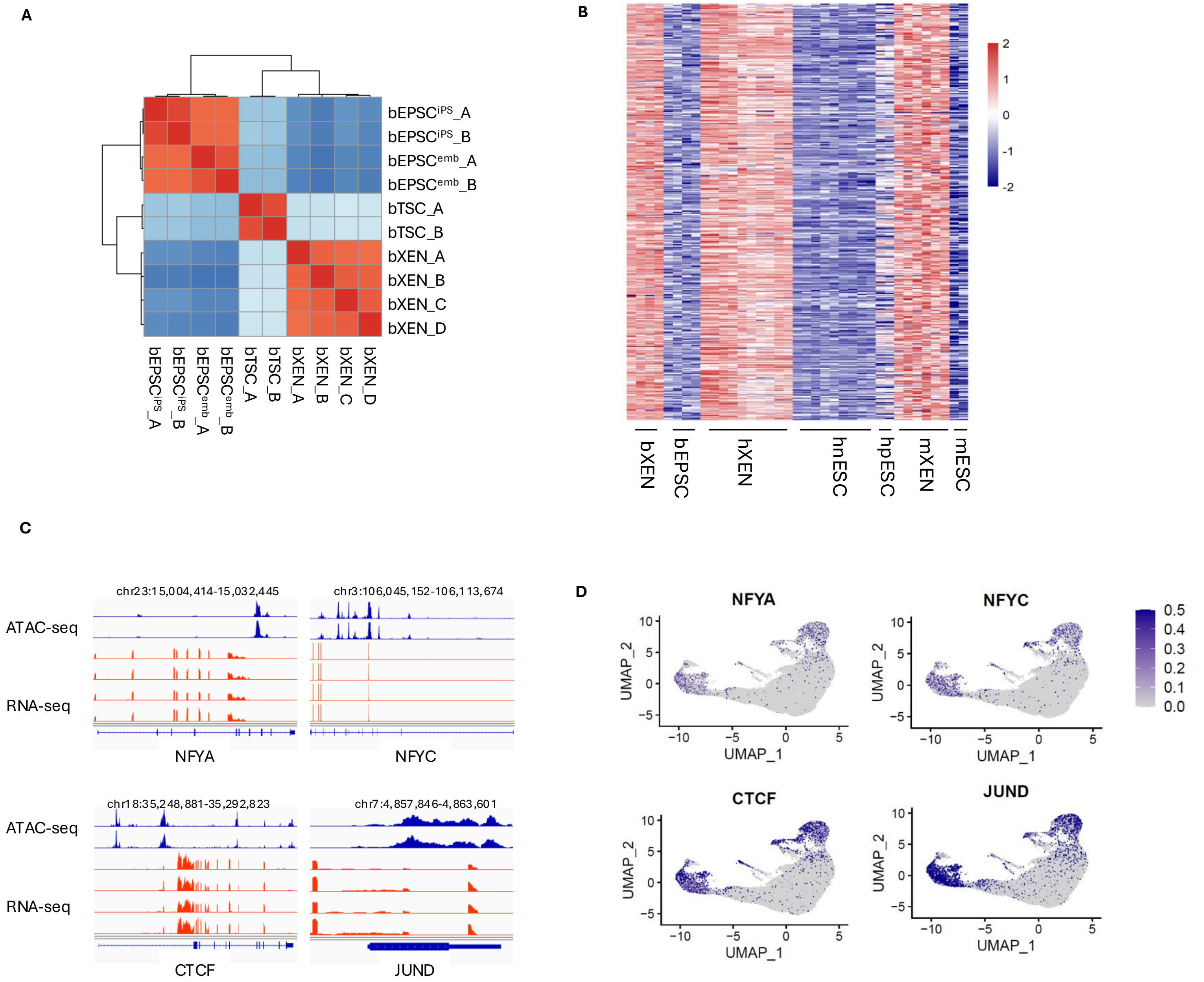

### Figure S3

*Figure S3*

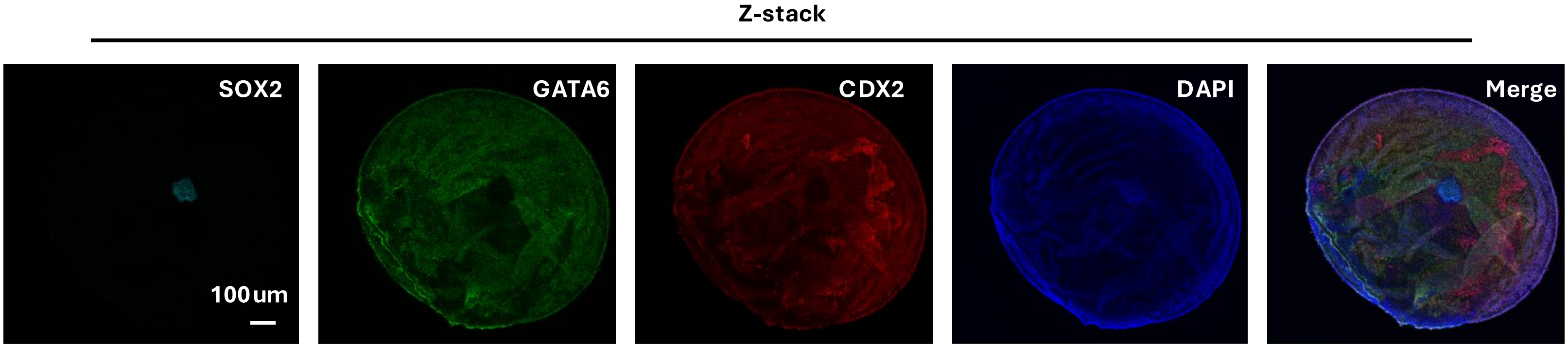

### Figure S4

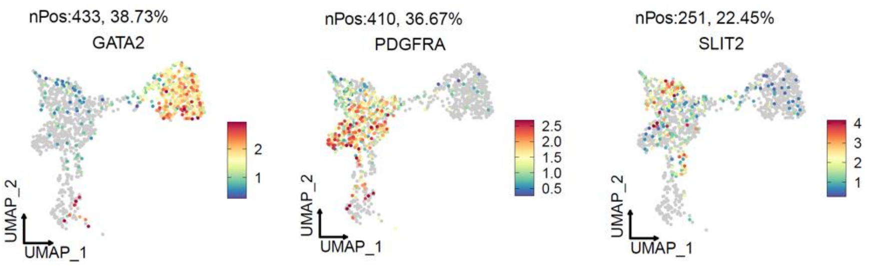

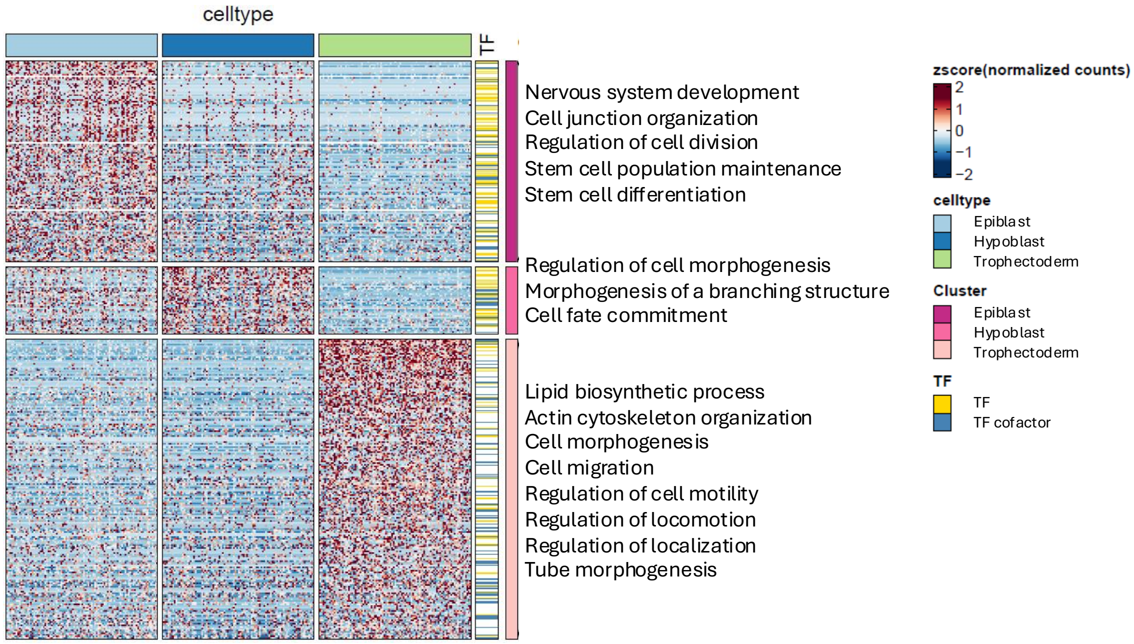
