## Supplementary material for "Establishment of bovine extraembryonic endoderm cells": Table S1

| | | M1 | M2 | M3 | M4 | M5 | M6 | M7 | M8 | M9 | M10 | M11 | M12 | M13 |
|---|---|---|---|---|---|---|---|---|---|---|---|---|---|---|
| **Basal medium** | N2B27 | ✓ | ✓ | ✓ | ✓ | ✓ | ✓ | ✓ | ✓ | ✓ | ✓ | ✓ | ✓ | ✓ |
| **Small molecules** | A83-01 | ✓ | ✓ | | | | | | ✓ | ✓ | ✓ | ✓ | ✓ | ✓ |
| | BMP4 | ✓ | ✓ | ✓ | | | | | ✓ | ✓ | ✓ | ✓ | ✓ | ✓ |
| | FGF4 | ✓ | ✓ | | ✓ | | | | ✓ | ✓ | ✓ | ✓ | ✓ | ✓ |
| | IL-6 | ✓ | ✓ | | | ✓ | | | ✓ | ✓ | ✓ | | ✓ | ✓ |
| | PDGF | ✓ | ✓ | | | | ✓ | | ✓ | ✓ | ✓ | | | |
| | XAV939 | ✓ | ✓ | | | | | | ✓ | ✓ | ✓ | | ✓ | ✓ |
| | retinoic acid | ✓ | ✓ | | | | | | | | | | | |
| | ActivinA | | ✓ | ✓ | ✓ | ✓ | ✓ | | ✓ | | ✓ | | | |
| | FGF2 | | ✓ | ✓ | ✓ | ✓ | ✓ | | | | | | | |
| | LIF | | ✓ | ✓ | ✓ | ✓ | ✓ | ✓ | | | | | | |
| | Chir99021 | | ✓ | ✓ | ✓ | ✓ | ✓ | ✓ | | | | | ✓ | |
| | IWR1 | | | | | | | | ✓ | | | | | |
| | DiM | | | | | | | ✓ | | | | | | |
| | MiH | | | | | | | ✓ | | | | | | |
| **Outgrowth rate (%)** | | 0/4 | 0/4 | 1/4 | 2/4 | 3/4 | 2/4 | 0/4 | 0/4 | 7/10 | 5/6 | 0/4 | 0/4 | 6/8 |
| **Long-term maintenance** | | | | | | | | | | ✓ | ✓ | | | ✓ |
